# Genomic Evidences Support an Independent History of Grapevine Domestication in the Levant

**DOI:** 10.1101/2020.07.11.198358

**Authors:** Aviad Sivan, Oshrit Rahimi, Mail Salmon-Divon, Ehud Weiss, Elyashiv Drori, Sariel Hübner

**Affiliations:** Galilee Research Institute (Migal), Tel-Hai Academic College, Upper Galilee 12210, Israel; Department of Chemical engineering, Ariel University, Ariel 40700, Israel; Department of Molecular Biology, Ariel University, Ariel, Israel; Department of Land of Israel Studies and Archaeology, Bar Ilan University, Israel; Eastern regional R&D Center, Ariel 40700, Israel

**Keywords:** grapevine, domestication, demography, transposable elements, whole-genome sequencing, selective sweep

## Abstract

The ancient grapevines of the Levant have inspired beliefs and rituals in human societies which are still practiced today in religious and traditional ceremonies around the world. Despite their importance, the original Levantine wine-grapes varieties were lost due to cultural turnovers commencing in the 7_th_ century CE, which lead to the collapse of a flourishing winemaking industry in this region. Recently, a comprehensive survey of feral grapevines was conducted in Israel in an attempt to identify local varieties, yet the origin of these domesticated accessions is unclear. Here we study the origin of Levantine grapevines using whole-genome sequence data generated for a diversity panel of wild and cultivated accessions. Comparison between Levantine and Eurasian grapevines indicated that the Levantine varieties represent a distinct lineage from the Eurasian varieties. Demographic models further supported this observation designating that domestication in the Caucasus region predated the emergence of the Levantine samples in circa 5000 years and that authentic descendants of ancient varieties are represented among the Levantine samples. We further explore the pedigree relationship among cultivated grapevines, identify footprints of selective sweeps, and estimate the extent of genetic load in each group. We conclude that the Levantine varieties are distinct from the Eurasian varieties and that resistance to disease and abiotic stress are key traits in the development of both Eurasian and Levantine varieties.

## Introduction

Since ancient times, domesticated grapevine (*Vitis vinifera* ssp. *sativa*) had an inspiring role in the culture of human societies. Among grapevine products, wine is the most popular and influential, thus many legends and believes were tied with its consumption in the Mediterranean region and the Near East (1,2). Today, the grapevine is among the most valuable horticulture crops in the world, cultivated on over 7 million ha. around the globe, mostly for wine production (3).

According to archaeological evidences, the grapevine was domesticated 8,000-10,000 years ago in the Taurus, Caucasus and Zagros Mountains (4). However, molecular evidences from chloroplast DNA suggest at least one more domestication event outside the Near East (5) albeit not conclusively (6). From the Caucasian region, domesticated grapevines have presumably distributed southwards to the Levant and later to Europe (1,4). Tracing the history of grapevines is challenging due to its spread as vegetative propagation material and mixtures between different genetic sources (2,7). Thus, the history of grapevine domestication and distribution remains largely obscure (8).

Recent genomic evidences allowed to study the domestication history of grapevine with higher confidence and to date the split between domesticated European varieties from wild grapevines to circa 22K years ago (9,10). These estimates significantly predate archaeological evidences, presumably due to a missing link to the direct wild ancestral population. Thus, the exact originating population of the domesticated grapevine is still a missing key in the domestication history of cultivated grapevines.

In the Levant region, winemaking was highly abundant at least since the Bronze age and has contributed significantly to the culture and economy of ancient societies. This important ancient industry has flourished for many centuries until the collapse of the Byzantine empire in the region during the 7_th_ century CE (11). For five centuries the wine industry was suppressed but continued to exist until the conquest of the Mamluk empire in the 13_th_ century when winemaking, cultivation, or production became completely forbidden (12,13).

Consequently, Levantine wine grapevine varieties which had an important role in ancient culture in this region for many centuries were abandoned and considered lost thereafter. Recently, a comprehensive grapevine survey was conducted in an attempt to revive the ancient Levantine wine industry (14). Based on a panel of SSR markers, the Levantine populations were clustered in a separate clade from European varieties (15). However, it is unclear whether the collected varieties are authentic ancient Levantine varieties or rather the outcome of a more recent introduction of European varieties during the 19_th_ century.

Here, we present the analysis of whole-genome sequencing data obtained for 81 domesticated (*sativa*) and wild (*sylvestris*) accessions representing a diversity panel of Levantine and Eurasian grapevines. We provide new evidences for the domestication history of grapevines in the Levant which support the authenticity of this material. Our results indicate that at least a few of the grapevine varieties that were cultivated in the Levant in ancient times survived the suppression of the wine industry in the region commencing in the 7_th_ century. In addition, genomic screening of the different populations provided evidences that the Levantine and Eurasian *sativa* lineages are distinguishable and that selective sweeps have affected known domestication syndrome traits in both groups in addition to group-specific loci.

## Results

### Population stratification in Eurasian and Levantine grapevines

A panel of 81 grapevine accessions was obtained from two independent sources. The Levantine panel included 46 putative domesticated *sativa* types and 9 wild *sylvestris* accessions (Fig. 1A) that were selected from a recently established collection genotyped previously with 22 SSR markers (15). Selected accessions were confirmed to be unique material based on the SSR markers and were whole-genome sequenced (WGS) to a coverage of 20X. In addition, publicly available raw WGS data for 17 Eurasian *sativa* varieties and 9 Eurasian *sylvestris* accessions were obtained from the SRA repository (9). Altogether 2.1 trillion bp were obtained for 81 grapevine accessions representing domesticated *sativa* and wild *sylvestris* types of Levantine and Eurasian origin. High quality trimmed reads from all accessions were aligned to the Pinot Noir PN40024 reference genome (16) followed by a variant calling procedure, yielding a total of 26,083,120 high quality (QUAL > 20) SNPs across all accessions. To further reduce the false positive rate among called variants, a machine-learning filtering approach was implemented and a set of 1,824,029 robustly called SNPs were kept for downstream analyses.

**Figure1.**
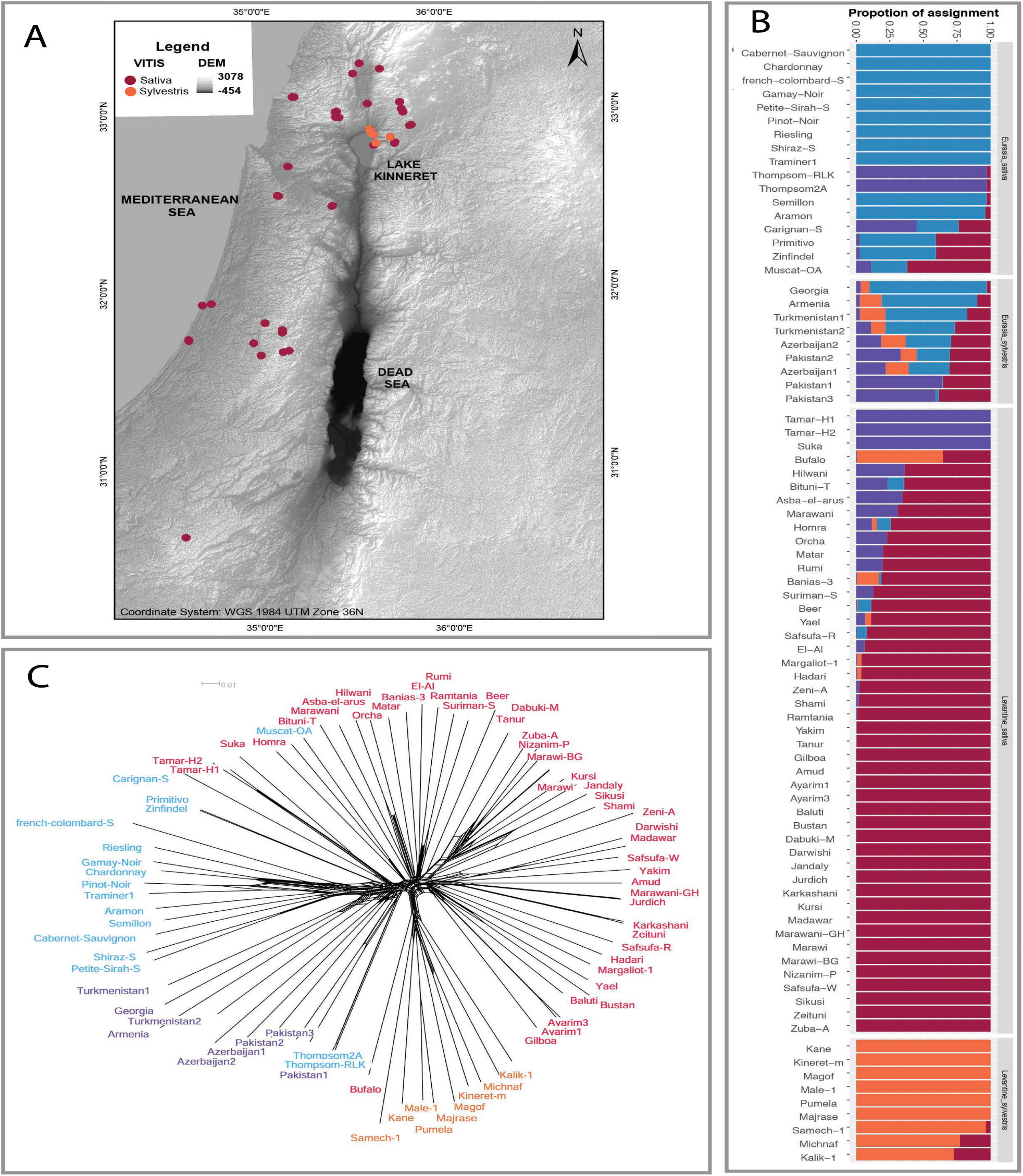
Population structure among all Eurasian and Levantine domesticated *sativa* and wild *sylvestris* grapevine accessions. **A)** Geographic map and locations where Levantine *sativa* (orange) and *sylvestris* (green) accessions were collected. **B)** Population structure among the 81 Eurasian and Levantine accessions. Analysis was conducted in FastSTRUCTURE and the barplot represents K = 4. Accessions are sorted by their expected group (top-bottom): domesticated Eurasian *sativa*, wild Euroasian *sylvestris*, domesticated Levantine *sativa* and wild Levantine *sylvestris*. **C)** Neighbor-joining network representing the resemblance between Eurasian *sativa* (red), Eurasia *sylvestris* (purple), Levantine *sativa*, and Levantine *sylvestris* (celeste) accessions. size

To investigate the population stratification among Levantine and Eurasian accessions, model-based analyses were conducted using fastStructure (17) (Fig. 1B, Fig. S1) and ADMIXTURE (Fig. S2) (18). Considering both analyses, the optimal clustering was obtained at K = 4 in accordance with the geographic origin and type of accessions: Levantine *sativa*, Levantine *sylvestris*, Eurasian *sativa*, and Eurasian *sylvestris*. Signs of admixture were observed among all four groups with the highest rate of admixed individuals in the Eurasian *sylvestris* group (100%), and lowest admixture rate in the Levantine *sylvestris* (33%). The difference in the level of admixture observed among the two *sylvestris* groups conceivably reflects the distribution range of each group, i.e. broad and narrow geographic range represented in the Eurasian and Levantine groups, respectively.

Overall, the assignment of accessions to clusters well supported previous characterization records with minor exceptions. Two accessions in the Eurasia *sativa* group (‘Thompson-RLK’, ‘Thompson2A’) known as table-grapes varieties were assigned to a distinct cluster from the remaining wine-grapes type. The same pattern was also observed in three Levantine *sativa* accessions (‘Tamar-H1’, ‘Tamar-H2’, and ‘Suka’) which were assigned to the same cluster as the two ‘Thompson’ varieties based on the fastStructure analysis. These results support previous reports for both Eurasian (9) and Levantine accessions (15). Strong signs of admixture were observed in specific accessions from both the Eurasian and Levantine *sativa* groups. For example, among the Levantine *sativa* accessions, a high rate of admixture with Levantine *sylvestris* (65%) was observed for the ‘Buffalo’ accession, presumably due to recent hybridization with wild Levantine grapevine (Fig.1 B). Among the Eurasian *sativa* accessions, high rates of admixture were observed in ‘Zinfandel’/’Primitivo’ (56%), ‘Muscat of Alexandria’ and ‘Carignan’ (74%).

To further investigate the level of divergence between the four grapevine groups, a neighbor-joining network was build using the SNP dataset called across all 81 accessions (Fig. 1C). The split into four groups in the network analysis supported the results of the model-based stratification analyses. Moreover, the obtained network clearly discriminated between the Eurasian and Levantine groups, implying that the Eurasian *sativa* group has branched from the Eurasian *sylvestris* group while the Levantine *sativa* group has branched from the Levantine *sylvestris* group. This pattern of divergence suggests that the Eurasian and Levantine *sativa* lineages do not share the same domestication history and may have developed in independent processes. To further confirm this observation, a PCA was conducted and designated the same pattern of divergence as obtained in the neighbor-joining network and model-based stratification analyses (Fig. S3). In addition, the network analysis well supported the mis-assignment of accessions identified by the model-based stratification analyses. Interestingly, one wild white-berry accession collected near the Sea of Galilee (‘Majrase’) was confirmed as a *sylvestris* type by all analyses. White-berry is a common phenotype among domesticated grapevines and considered a post-domestication characteristic (19). A white-berry phenotype in wild grapevine is possibly the result of introgression from cultivated *sativa*, however, none of the analyses conducted supported signs of introgression from cultivated grapevines into this wild white-berry accession.

Next, the level of nucleotide diversity (π), Watterson’s θ, Tajima’s D, observed heterozygosity, and linkage disequilibrium (LD) were investigated in each group after excluding five mis-assigned accessions (defined as less than 10% assignment to their expected cluster). Linkage disequilibrium analysis conducted across all *Vitis* accessions indicated that decay is reached at 20Kbp (Fig. S4, Table S1). Within groups, steep LD decay and high genetic diversity were observed among the Eurasian *Sylvestris* (π = 0.29 ± 0.01, LD= 8 Kbp) compared with the Eurasian *sativa* group (π = 0.27 ± 0.01, LD = 10 Kbp) which is characterized by a significantly lower diversity (*t* = 3.50, *p* =1.20×10 _-4_) due to domestication bottleneck (Table S1). An opposing trend was observed among the Levantine groups, i.e. steep LD decay and high genetic diversity within the Levantine *sativa* group (π = 0.27 ± 0.01, LD = 6 Kbp) compared with Levantine *Sylvestris* (π = 0.25 ± 0.01, LD = 25 Kbp) which was also characterized by a significantly lower diversity (*t* = 6.35, *p* = 2.30×10_-7_). This pattern among the Levantine groups is attributed to the constrained geographic range of wild grapevines at its southern distribution edge (15). In addition, both the population stratification and genetic diversity analyses imply that the Levantine *sativa* group is likely a mixture of two distinct subgroups (Fig 1B). Thus, the domesticated Levantine *sativa* group was further split into two subgroups (Table S1), a homogenous Levantine *sativa*, and a highly diverse group with signs of admixture with other genetic sources.

### The demographic history of Eurasian and Levantine grapevines

To test the hypothesis that Eurasian and Levantine *sativa* are distinct lineages that were developed in two independent processes, various complementary demographic analyses were conducted. To reduce the confounding effect of differences in sample size and level of admixture across groups, nine representative accessions were selected from each group to adjust the sample size to the smallest groups (nine accessions in Eurasia *sylvestris* and Levantine *sylvestris*). To explore historical splits and gene-flow among the four groups, a graphical model-based analysis was conducted using TreeMix (20). The SNP dataset was restricted to intergenic regions to reduce the effect of selective sweeps on the demographic inferences, thus a total of 1,055,512 SNPs was kept for the analysis. The model uses the allele frequencies in modern populations and a Gaussian approximation to infer historical demographic events. First, the model was carried out without migration (*f*_index_ = 0.996) and the obtained graph supported the hypothesis of independent demographic history (Fig. 2A). Although the model does not allow to precisely estimate times of split events, it was noted that domestication in Eurasia predated the development of the Levantine *sativa*. Allowing one migration event in the model improved its likelihood (*f*_index_ = 0.999) and indicated a historical gene-flow from Levantine *sativa* into the Eurasia *sativa* group (Fig. 2B). This inference supports previous reports on the possible exchange of germplasm by the Romans, crusaders, or Islamic rulers (2). Allowing more migration events in the model did not improve its likelihood (Fig. S5).

**Figure2.**
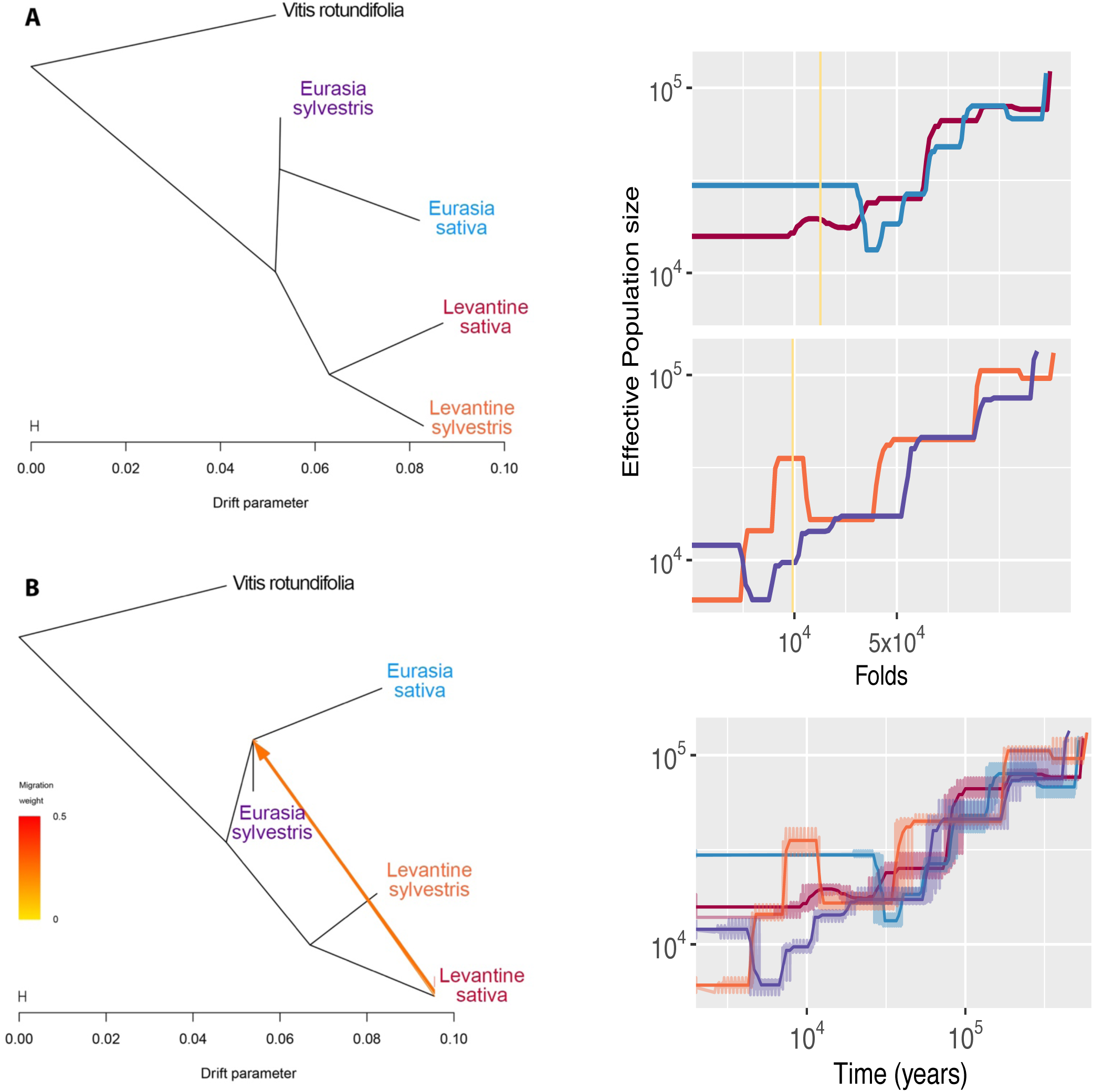
Demographic analysis of nine representative accessions of each of the four grapevine groups: Eurasian *sativa* (blue), Eurasian *sylvestris* (purple), Levantine *sativa* (red) and Levantine *sylvestris* (orange). **A+B)** Inference of splits and gene-flow among the four grapevine groups. The analysis was conducted in TreeMix using *Vitis rotundifolia* as an outgroup to root the tree. Presented are the results for the analysis without migration (A) and with one migration event (B). When migration was allowed in the model, direction of the migration is indicated with arrow and its color represent the migration weight. **C)** Demographic history inference among the four grapevine groups. Analysis was conducted with SMC++ for all four groups without the ‘clean split’ model (bottom) and for the Eurasian *sylvestris-sativa* pair and Levantine *sylvestris-sativa* pair with the ‘clean split’ model (top). The estimated split time in the models is indicated with yellow vertical line. The x-axes represent time in years where the leftmost part correspond to recent time. The y-axes correspond the estimated effective population size. Each SMC++ analysis was conducted with 10 cross-validations procedures and 20 iterations. Confidence intervals are indicated with light colors in the plot.

To further investigate the demographic history of grapevines in Eurasia and the Levant, a coalescence analysis was conducted using the SMC++ program (21) which allows to incorporate information from all nine individuals representing each group. To infer the demographic changes on a timescale, a generation time of three years and a mutation rate of 5.4×10 _-9_ per nucleotide per generation were used in the model (Fig. 2C).

The SMC++ analysis was conducted and inspected with and without the ‘clean-split’ model to allow a better interpretation of the demographic process. In all models, the divergence between groups predated the ‘clean-split’ model indicating that grapevine domestication was prolonged over a period of time until the two groups were fixed as distinct lineages. In agreement with previous studies (9,10), the SMC++ model denoted that the divergence between Eurasian *sylvestris* and *sativa* grapevines commenced circa 30,000 years ago and reached a clear differentiation about 15,000 years ago based on the ‘clean-split’ model (Fig. 2C). Conducting the analysis for the Levantine *sativa* group indicated that divergence from Eurasian *sylvestris* and *sativa* predates the estimated domestication event in Eurasia (20,000 and 17,000 years ago, respectively). The same demographic analysis between Levantine *sylvestris* and *sativa* indicated that divergence between these groups commenced approximately 15,000 years ago but they reached a clear differentiation 10,000 years ago according to the ‘clean-split’ model. Considering the small distribution range represented among Levantine *sylvestris* accessions and the long period of divergence, it is unlikely that the wild population represented in this study is the direct progenitor of Levantine *sativa*. Nevertheless, the demographic inferences for the Levantine *sativa* group provide evidence that these accessions are authentic Levantine varieties that sustained the suppression of grapevines cultivation and winemaking industry in the Levant. Arguably, the descendants of the lost Levantine wine varieties that were cultivated in this region in ancient times. To further validate these results, a second analysis was conducted with the MSMC software (22) using four individuals that were randomly sampled from each population. The obtained trajectories in the effective population size of each group over time supported the observed split between the Eurasian *sylvestris* and *sativa* grapevines circa 20,000 years ago and a more recent split between the Levantine *sylvestris* and *sativa* grapevines circa 9,000 years ago. The MSMC inferences accord with the results obtained by the SMC++ and TreeMix analyses (Fig. S6).

### Pedigree network in domesticated Eurasian and Levantine varieties

To better understand the recent history and pedigree relationship among grapevine varieties, a relatedness network was constructed for all 58 domesticated (*sativa*) accessions of both Levantine and Eurasian origin. The relatedness matrix was computed at 1cM segments using the refined-IBD tool (23) which allows to identify traces of shared ancestry signatures. This approach has the advantage of distinguishing between ‘old’ ancestry (short segments) and recent ancestry (long segments) signatures. The minimum threshold to delineate a parent-offspring or sibling relationship was determined based on a confirmed cutoff (IBD = 0.466) which was computed using a short segments ancestry detection approach (7). To link between this confirmed cutoff and the calculated refined-IBD score, a correlation analysis was conducted between overlapping pairs of accessions present in both studies (spearman: *rho* = 0.62, *p* = 0.028). To graphically visualize the pedigree network, a first-degree relationship graph was constructed using Gephi (24) which allows organizing accessions in an attraction-repulsion network (Fig. 3). Overall, the Levantine *sativa* formed a distinguished cluster from the Eurasian *sativa* group which further support the independent history of the Levantine domesticated accessions. Clonal relationships were identified in both the Eurasian (‘Zinfandel’/‘Primitivo’) and Levantine (‘Zeni-A’/’Suriman-S’/’Asba-el-arus’) groups. Interestingly, a pedigree relationship was observed between the European ‘Chardonnay’ variety and Levantine accessions sampled at the Galilee (‘Margaliot-1’, ‘Hadari’).

**Figure3.**
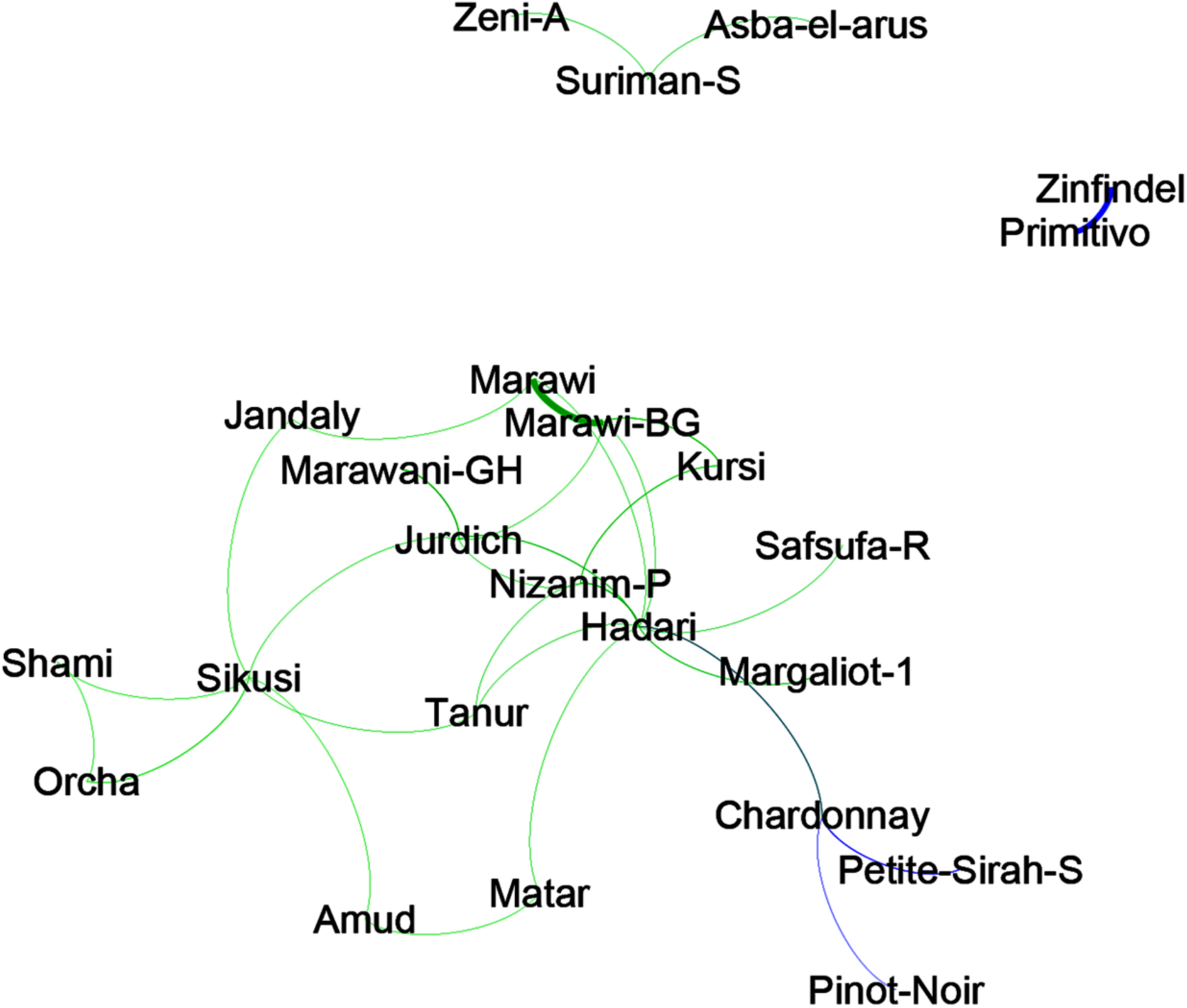
Relatedness among domesticated Eurasian and Levantine *sativa* accessions. Relatedness was calculated from identity by descent (IBD) in 1cM fragments across all 57 *sativa* accessions. In the network, only strong links higher than 0.466 are indicated and the thickness of the line represents the link strength. Links between Levantine accessions is indicated with green edges and links between Eurasian accessions are indicated in blue edges.

### Footprints of domestication in cultivated verities

To investigate whether the independent history of the Levantine and Eurasia *sativa* has resulted in distinct patterns of selective sweeps we performed a genome scan analysis using the *µ*-statistic score (25) that was calculated for the 9 representative accessions from each group. The *µ*-statistic is a composite score combining the site frequency spectrum (SFS), linkage disequilibrium, and genetic diversity calculated at overlapping windows that are adaptively determined according to the calculated metrics. We considered a significant outlier window following the default cutoff of 99.95% (Fig. 4). Altogether, 5,581 outlier windows were detected in the Eurasia *sativa* group and 6,333 outlier windows were detected in the Levantine *sativa*. The average *µ*-statistic in the Levantine *sativa* group was significantly higher than in the Eurasian *sativa* group (*t* = 10.6, *p* < 0.0001). The overall stronger signal of selective sweep detected in the Levantine *sativa* group may indicate that this group has experienced stronger selection or that it was developed more recently. To identify footprints of a selective sweep as a result of domestication and breeding, overlapping windows were merged and extended to form quantitative trait regions (QTRs). Merging was performed only for outlier windows found in distance shorter than 20Kbp in accordance with LD decay in *vitis*. A total of 222 and 260 outlier QTRs were detected across all chromosomes in the Eurasian and Levantine *sativa* groups, respectively (Table S2). Among the identified QTRs, 80 were found in both groups across all chromosomes except for chromosome 8 where no QTR was detected in neither group. Enrichment in the number of QTRs was observed in the Eurasian *sativa* group on chromosomes 12 (26 QTRs), 13 (17 QTRs), and 19 (22 QTRs), and a strong signal of selection (*µ* > 100) was observed on chromosomes 7 (103.1) and 13 (145.5). In the Levantine *sativa* group, enrichment in the number of QTRs was observed on chromosomes 4 (25 QTRs), 5 (32 QTRs), and 18 (21 QTRs), and a strong signal of selection (*µ* > 100) was observed on chromosomes 2 (110.8), 11 (751.5), 12 (106), 17 (409.8), and 18 (150.4).

**Figure 4.**
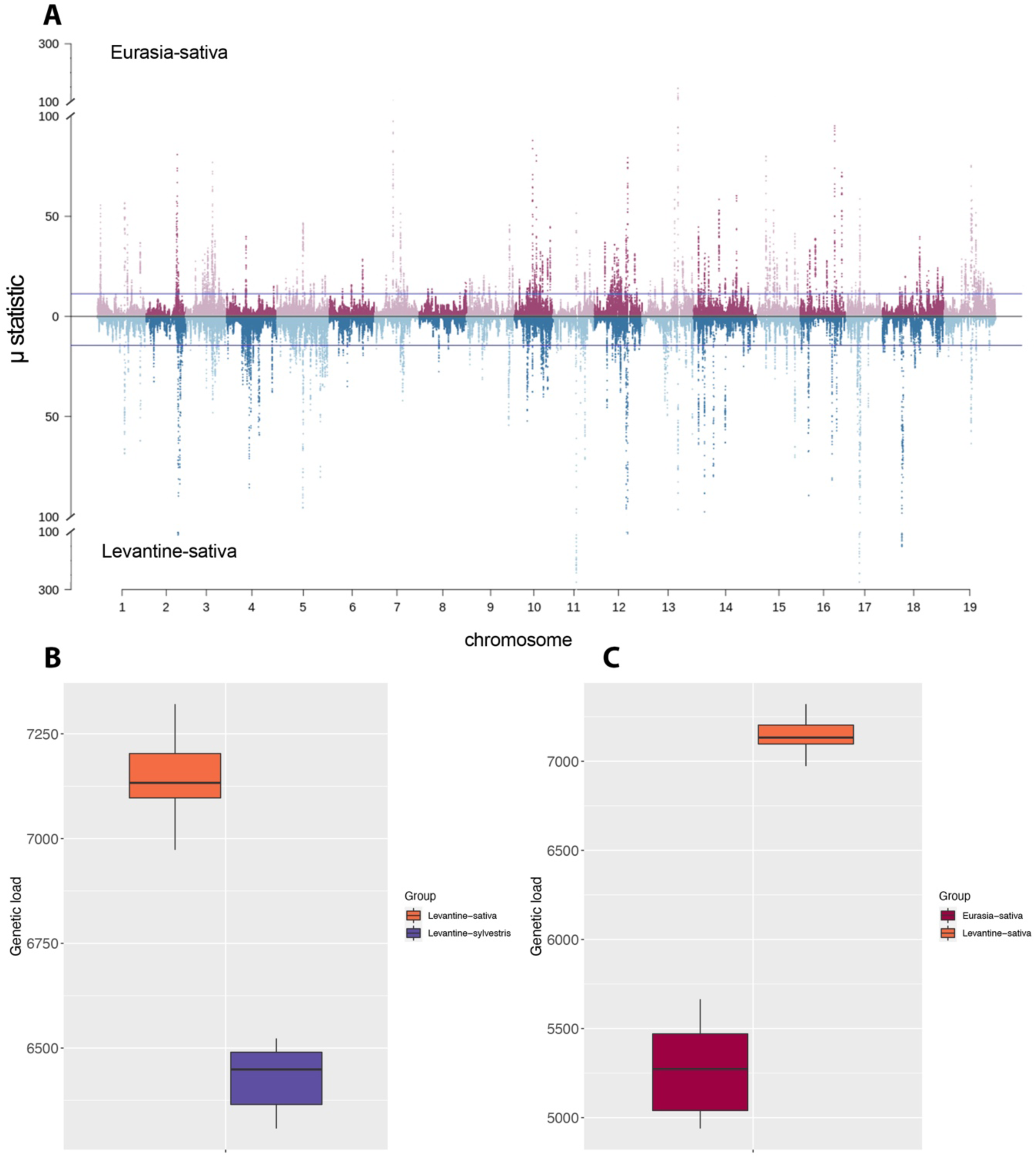
**A)** Signatures of selective sweeps in Levantine and Eurasian grapevines. Selection signal was measured **using** the μ-statistic in sliding windows. Each dot represents a SNP score in the Levantine *sativa* (blue) and Eurasian *sativa* (red). **B)** Comparison between genetic load calculated in Levantine *sativa* and Levantine *sylvestris* groups. C) Comparison between genetic load calculated in Levantine *sativa* and Eurasian *sativa* groups.

Several candidate genes were found within outlier QTRs in both Eurasian and Levantine *sativa*. For example, the genes resveratrol synthase and stilbene synthase that are involved in the response to biotic stress as well as in taste and aroma (26) were found in an overlapping QTR on chromosome 16 in both groups. Another example was observed on chromosome 2 in both *sativa* groups where bifunctional nitrilase/nitrile hydratase genes were found. These genes were recently targeted for their potential role in grapevine domestication (9,27). Among the Levantine *sativa* group, a strong signal of a selective sweep was observed on chromosome 18 where a cluster of phenylalanine ammonia-lyase (PAL) genes was identified. PAL genes contribute to anthocyanin concentration in the berry pericarp which affects berry color and wine quality in addition to enhancing resistance to biotic and abiotic stresses (28). In the same QTR located on chromosome 18, we also detected an RPW8 gene that confers a basal resistance to powdery mildew in Arabidopsis (29). Other QTRs that were detected exclusively in the Levantine *sativa* group were on chromosome 17 where several abiotic stress-responsive genes were identified including the basic helix–loop–helix transcription factor (30) and HVA22-like gene (31,32).

Outlier QTRs that were detected exclusively in the Eurasian *sativa* group harbored the anthocyanin synthesis genes (MYBA1 and MYBA3) on chromosome 2 (32,33), the pathogen response gene (ACD6) on chromosome 16 (35), and the disease response genes (RPM1) on chromosome 7 which were described to be up-regulated in response to pathogen infection in grapevine (36).

### Genetic load among cultivated grapevines

To evaluate the extent of genetic load due to the accumulation of deleterious mutations along the genome in each of the *sativa* groups we used the SIFT4G software (37). A total of 92,936 non-synonymous variants were detected of which 37,635 were predicted as deleterious (SIFT score < 0.05). To correct for potential bias introduced by the use of a reference genome, alleles identified also in the outgroup species *V. rotundifolia* were not considered deleterious (9). Altogether 29,386 sites were variants were identified as deleterious across all accessions and used to calculate the genetic load in each group. Not surprisingly, the calculated genetic load in the Levantine *sativa* group (7,142 α 101) was significantly higher (*t* = 4.51 *p* = 1.70×10_--4_) than in the Levantine *sylvestris* group (6,228 α 598) due to the effect of domestication. Moreover, significantly higher genetic load (*t* = 14.70, *p* = 1.24×10_-4_) was observed in the Levantine *sativa* group compared with the Eurasian *sativa* (5,168 α 390). This observation supports the results obtained in the selective sweep analysis indicating that the Levantine group was under stronger selective pressure or was generated more recently.

## Discussion

Along the history of human societies, wine has provided a special cultural flavor to the life of ancient and modern societies. According to archaeological observations, the grapevine was domesticated and spread by ancient societies in the Near East (circa 10,000 BC), and later was introduced to East Asia, the Mediterranean basin and Europe (4,10).

In the Levant region, grapevine cultivation has flourished for several millennia until the collapse of the Byzantine empire during the 7_th_century CE (11). Since then, wine production has declined and eventually abandoned under the Mamluk empire conquest, and the ancient grapevine varieties were considered lost (12,13). In this study, we provide new genomic evidences for the demographic history of grapevine varieties in the Levant, their origin, and the genomic landscape of their domestication.

### The origin of cultivated grapevines in the Levant

To study the history of Levantine varieties we compared whole-genome sequence data from *sylvestris* and *sativa* types of Levantine and Eurasian origin. All population stratification analyses supported the deviation into four distinct groups by type (*sylvestris*/*sativa*), and geography (Levant/Eurasia). Moreover, the clustering pattern implied that the Eurasian *sativa* group has branched from the Eurasian *sylvestris* group while the Levantine *sativa* group branched from the Levantine *sylvestris* group, with few minor exceptions (Fig. 1). In accordance with previous studies (9), the Eurasian table-grapes varieties (‘Thompson’) and ‘Muscat of Alexandria’ are distinguished from remaining varieties and a similar pattern was also observed among three Levantine *sativa* accessions which presumably represent introduced table-grapes varieties (‘Tamar-H1’, ‘Tamar-H2’, ‘Suka’). Also, one Levantine accession was identified as a potential feral grapevine (‘Buffalo’), and one white-berry accession (‘Majrase’) was confirmed to be of wild origin. The latest could be an interesting example of a sporadic occurrence of white-berry mutation in the wild, as we failed to identify signs of admixture with other domesticated varieties.

The process of domestication can be generally divided into three phases which include management of wild material, selection of desirable basic domestication traits, and dispersal of the domesticated material. During the third stage of domestication, introgression from local wild populations can increase adaptation of the alien crop to the local environment (8,38). These introgressions may be spread across the genome yet the genetic background should reflect the origin of the crop. Once the genetic turnover is so profound that the origin is masked by introgression, the question of the origin of domesticates becomes quantitative. The demographic analyses conducted in this study supported an independent domestication history of the Levantine *sativa* since no traces of late introgressions were identified. However, we cannot exclude based on our analyses a complete genomic turnover of the original domesticated material in the Near East. In accordance with previous studies (9,10), our analyses pointed that the divergence of Eurasian *sativa* from *sylvestris* commenced approximately 30 thousand years ago, however, the ‘clean-split’ model in SMC++ provided an improved estimate for the domestication of grapevine in Eurasia to approximately 15,000 years ago. It should be noted that these models are limited in their ability to estimate recent demographic events especially in the past few centuries (21). Therefore, it is difficult to infer robustly from these models whether the Levantine accessions are truly descendants of ancient varieties or the outcome of a recent introduction of grapevine varieties from Eurasia. The pedigree and genome scan analyses imply these are indeed authentic Levantine varieties.

### Pedigree relationship between Eurasian and Levantine varieties

To test for potential recent admixture between Levantine and Eurasian *sativa*, and the effect of vegetative propagation on the similarity between cultivars (7), a relatedness network was constructed based on pair-wise identity by descent (IBD) analysis. In accordance with previous studies (7) conducted for the Eurasia *sativa*, clonal relationship was observed among the Eurasian accessions and also among the Levantine accessions but no pedigree links were observed between the two groups with one exception. A pedigree link between the European ‘Chardonnay’ variety and Levantine accessions sampled at the Galilee region (‘Margaliot-1’, ‘Hadari’) imply a potential Levantine ancestry for ‘Chardonnay’. Previously, ampelographers suggested that ‘Chardonnay’ has ancestral roots in the Levant, although it was later contradicted by genetic analysis which indicated that ‘Chardonnay’ was produced by a cross between ‘Pinot Noir’ and ‘Gouais blanc’ (39,40). While ‘Pinot Noir’ is a confirmed French variety, ‘Gouais blanc’ is considered an introduced variety from elsewhere. The results obtained from the pedigree analysis do not allow to track back the entire lineage of ‘Chardonnay’ and how it is linked to the Levant, but to the best of our knowledge, this is the first genomic evidence for this ancestry relationship.

### Development of grapevine varieties in Eurasia and the Levant

All population stratification, demography, and pedigree analyses conducted supported the hypothesis that Levantine *sativa* originated from a different source than the Eurasian *sativa*. To test whether these independent histories are reflected in different genomic footprints of domestication, genome scans were conducted for each *sativa* group. Footprints of selective sweeps were detected across all chromosomes in both groups with many overlapping QTRs. However, several group-specific QTRs were also detected and included candidate genes that are involved in biotic and abiotic stress resistance. Environmental stress has always troubled farmers and breeders around the world, however, since the pathogen identity and type of stress varies between regions, so are the resistance genes selected to confront them.

The revolution of genomics provides a powerful tool to fill gaps in the history of crops at multiple genomic levels (41). Despite their importance in human culture and economy, many ancient Levantine grapevines varieties were consider lost for many centuries. Here we provided evidences that the Levantine ancient grapevine varieties survived, in some cases under harsh conditions, despite the cultural turnovers in this region along history. Nevertheless, to link between the discovered Levantine accessions and ancient varieties we need to obtain high-quality sequence data from archaeological samples which is now in reach also for grapevines.

## Materials and methods

### Whole-genome sequencing and variant calling

Samples were obtained from two separate sources (Table S3). Levantine samples were selected from a recently established large *Vitis* collection comprised of 372 accessions which were genotyped using 22 standard SSR markers (15). Based on this data, clone accessions were removed and a representative diversity panel of 55 accessions was selected for our study including 46 accessions identified as *sativa* and 9 accessions identified as *sylvestris*. From each accession, a young leaf tissue (shoot tip) was sampled for DNA extraction using the QIAmp DNA Micro cleanup kit (Qiagen, Valencia, USA), and a library was prepared using the NEBNext Ultra DNA library preparation kit (catalog number E7370L; NEB). Whole-genome sequencing of 150bp paired-end reads was generated on Illumina HiSeq2500 machine to a target coverage of 20X per sample. Raw sequence data for 17 Eurasian *sativa* and 9 *sylvestris* samples were obtained from the short-read archive (SRA) project number PRJNA388292 (9). Altogether, whole-genome sequence data of 81 accessions were obtained and analyzed.

The quality of raw sequence data for all Levantine and Eurasian samples was inspected using FastQC v0.11.8 (42). Low-quality reads were trimmed and adapters were removed using Trimmomatic v.0.32 (43) with default parameters. Following trimming, reads were inspected again with FastQC to guarantee only high-quality reads are included in the analysis. Clean reads from each sample were aligned to the Pinot Noir 40024 reference genome (16) using BWA-MEM (44) with default parameters. The obtained alignment files were further processed following the GATK best practice protocol (45) and included marking duplicates with picard-tools v.2.8.1 (46), realignment around indels with GATK (47) and indexing using samtools v.1.3.1 (48). Variant calling was conducted across all 81 accessions in one batch using the HaplotypeCaller program and variants were filtered using the variant quality score recalibration (VQSR) algorithm as implemented in GATK v3.6 (47). Briefly, the VQSR uses a machine-learning algorithm to develop a model of true variants based on validated SNPs and allows the discrimination between true and false calls. We used the 20K Illumina SNP-chip data (49) as a training set in the VQSR analysis and a minimum LOD score of 4 was set to guarantee that the highest confident SNPs are kept for downstream analyses. The obtained SNP set was further filtered to exclude sites with more than 20% missing data across all samples and a minimum minor allele frequency of 5%.

### Population stratification analyses

For population structure analysis we used both the FastStructure v1.0 (17) and ADMIXURE v1.3.0 (18) programs. For each analysis, the number of ancestral populations (K) tested ranged from 2 to10 with 20 replicates for each K. The cross-validation procedure implemented in ADMIXTURE and the ‘chooseK’ tool implemented in FastStructure were used to select the most likely number of clusters explaining the population structure among *vitis* accessions.

A neighbor-joining (NJ) network was constructed in SpitsTrees4 (50) using all SNPs that passed the filtering procedure. In addition, a principal component analysis (PCA) was conducted using the smartPCA program as implemented in EIGNSOFT v6.1.4 package (51).

### Demographic analyses

To infer the historical relationship including events of splits and migrations among populations, we used the TreeMix v1.13 program (20). To avoid sample size bias on demographic inferences the number of accessions in each group was set to nine individuals in accordance with the smallest groups (*sylvestris* groups). In each *sativa* group, the 9 accessions with the highest ancestry assignment, as obtained by FastStructure, were selected for downstream analyses. To avoid bias introduced by genomic regions affected by non-neutral processes, the SNP dataset was restricted to intergenic regions. In addition, SNPs were called in a *Vitis rotundifolia* accession (Muscadine, SRA accession number: SRR5627788) (9) which was used in the model as an outgroup for rooting the tree. To filter non-independent SNPs in the model, windows were set to the size of twice the calculated average number of SNPs per 20Kbp (the evaluated extent of LD across populations). Additionally, zero to four migration events were tested in the model.

To further infer the demographic history and split time in the Levantine and Eurasian populations, we used the likelihood-free SMC++ program v1.15.2 (21). The SMC++ program allows to leverage information from multiple individuals of each population and infer changes in the effective population size also in recent history. A mutation rate of *µ* = 5.4×10 ^-9^ mutations per base-pair per generation, and a generation time of 3 years were assumed in all models. The input data from each population was filtered for long stretches of homozygosity (>20Kbp) and the cross-validation (CV) module was used to infer the effective population sizes with 10-fold CV steps. A polarization error rate was set to 0.5 to allow uncertainty on the identity of the ancestral allele. Finally, the ‘clean-split’ model was used with default parameters to estimate split times between pairs of populations.

In addition, the MSMC v2 program (22) was used to estimate changes in the effective population size (*N*_e_) in each population over time. Four individuals were randomly chosen from each population after phasing and imputing the entire SNP data set using BEAGLE v5.1 (39). A mutation rate of *µ* = 5.4×10^−9^ mutations per base-pair per generation and a generation time of three years were set in the model.

### Pedigree network analysis

To explore the breeding history of grapevine verities in the Levant and their relationship to the Eurasian varieties, a pair-wise identity by descent (IBD) was calculated across all domesticated *sativa* accessions (Eurasia and Levant). The Refined-IBD program in BEAGLE v5.1 (23) was used to calculate the pair-wise IBD, and the results were converted to a kinship score using the ‘relatedness’ tool as implemented in BEAGLE v5.1 (39). A relatedness threshold of 0.466 was set following a previous study (7) on the familial relationship among grapevine varieties based on IBD scores calculated in the program plink (52). To adjust the relatedness scores calculated from the Refined-IBD to the pre-calculated threshold, a Spearman correlation analysis was conducted between the Refined-IBD and the plink scores computed for each pair of accessions. The obtained pair-wise IBD matrix was visualized with the network analysis program Gephi v0.9.2 (24) using the multi-gravity force-atlas2 algorithm. This algorithm allows to transpose the IBD kinship in the attraction-repulsion 2D network and the sub-networks were identified using the modularity algorithm which considers both quality and quantity of the pair-wise links.

### Population genomics statistics

Linkage disequilibrium (LD) was computed using the PopLDdecay software (53) across all accessions and for each population separately. LD was also measured within each population at 1Mbp windows using plink (52). Population diversity statistics were calculated for each group separately using the PopGenome package (54) and included nucleotide diversity (π), Tajima’s D, and Watterson’s θ.Observed heterozygosity was obtained for each population using VCFtools v0.1.15 (55).

Genome scans for the footprint of selective sweeps were conducted within each domesticated *sativa* group. Within each group, the *µ*-statistic was calculated using the RAiSD software for 9 accessions from each group (25). The *µ*-statistic is a composite score of the changes in site frequency spectrum (SFS), linkage disequilibrium, and genetic diversity. Top-ranked windows (>99.95%) were considered outliers. Overlapping windows with a maximum gap of 20Kbp were merged with BEDtools v2.26.0 (56) to allow a comparison of genomic regions between groups.

### Genetic load estimation

To estimate the genetic load in each accession we first identified non-synonymous mutations using the *Vitis vinifera* reference genome (NCBI_Assembly: GCF_000003745.3) and associated annotation files (annotations release 102, GCF_000003745.3_12X) to build a SIFT genomic database in SIFT4G (37). The identified non-synonymous mutations within coding regions were considered as potentially deleterious if the obtained SIFT score was lower than 0.05. To avoid a reference bias effect on the genetic load predictions, alleles identified also in the outgroup species *Vitis rotundifolia* were not considered deleterious. The genetic load was calculated by summing the number of deleterious alleles in each accession with a score of one for heterozygote and two for homozygote deleterious alleles.

## Acknowledgment

This work was supported by the JNF grant # 90-23-020-12, and the general support of the Israeli Ministry of Science and Technology (MOST).

## Author contribution

ED, EW, and SH designed this study; AS performed the research; AS, OR and ED analyzed the data; and AS, ED, EW, and SH wrote the manuscript.

